# PN6047 Demonstrates Broad Preclinical Efficacy in Headache Models as a Novel Delta-Opioid Receptor Agonist

**DOI:** 10.64898/2026.06.09.729278

**Authors:** Yaseen Awad-Igbaria, Yanping Zhang, Mattin Aframian, Guido C. Faas, Andrew Charles, Serapio M. Baca, Emily Jutkiewicz, Bengt von Mentzer, John Traynor, David Kendall, Amynah A. Pradhan

## Abstract

**Background:** The Delta-opioid receptor (DOR) has gained attention as a promising target for the treatment of migraine and headache disorders. This is largely attributed to its unique pharmacological profile, which suggests that DOR-targeting treatment offers effective therapeutic benefit with a lower risk of medication overuse headache (MOH), reduced abuse liability, and minimal potential for physical dependence. These advantages have driven the development of a novel DOR agonist PN6047 (3-[[4-(dimethylcarbamoyl) phenyl]-[1-(thiazol-5-ylmethyl)-4-piperidylidene] methyl]benzamide), which has completed Phase I clinical trial and showed a favorable safety and tolerability profile. Although PN6047 has shown promising effects in neuropathic pain models, its efficacy in preclinical models of headache-associated pain remains to be evaluated. Here, we investigated the effects of PN6047 in models of migraine-associated pain and aura as well as post-traumatic headache (PTH) and MOH.

**Methods:** C57BL6/J mice were used to examine the effects of PN6047 in the following migraine models: chronic intermittent nitroglycerin (NTG)-induced migraine-associated pain, PTH, KCl-induced cortical spreading depression (CSD), and optogenetic evoked CSD in a freely behaving transgenic mice expressing ChR2-eYFP. In addition, we tested whether chronic PN6047 induced MOH and whether it could prevent the development of MOH induced by sumatriptan.

**Results:** A single injection of PN6047 blocked chronic cephalic allodynia established by chronic intermittent NTG and PTH. Moreover, chronic PN6047 treatment prevented the development of MOH induced by sumatriptan, without causing MOH itself. In addition, PN6047 significantly reduced the number of CSD events in the KCl-induced CSD model, and delayed CSD onset triggered in freely behaving mice along with subsequent CSD-evoked allodynia.

**Conclusion:** PN6047, a novel DOR agonist, strikingly blocks headache-associated mechanism and symptoms in preclinical models of chronic migraine, migraine aura, PTH, and MOH. Importantly, prolonged PN6047 treatment did not induce MOH or analgesic tolerance. Together, these data demonstrate that despite the distinct mechanisms underlying migraine and headache disorder, PN6047 exhibits robust efficacy without inducing MOH, and displays a favorable safety and tolerability profile.

## Introduction

Headache disorders are among the most prevalent causes of years lived with disability worldwide (1, 2), affecting approximately 3 billion individuals globally (3). Among headache disorders, migraine accounts for most of the overall burden (2, 4). Chronic migraine is recognized as the most disabling form of migraine, which is defined as 15 or more headache days per month lasting for at least 3 months (5), and affects 1-2% of people in US (6). Interestingly, longitudinal studies indicate that about 2.5% of patients transition from episodic to chronic migraine yearly (7). One of the risk factors that contributes to chronic migraine progression is overuse of acute migraine therapies such as triptans, barbiturates, or opioids (8, 9). Notably, medication-overuse headache (MOH) has been observed in a substantial proportion of patients with chronic migraine (8, 9). Beyond the risk for MOH of migraine therapies, a large proportion of migraine patients are still not fully satisfied with their treatment (10, 11). These challenges drive the need for the therapeutic development of safe and more effective therapeutic strategies.

The delta opioid receptor (DOR) has gained attention as a novel and promising target for the treatment of migraine, headache disorders and chronic pain (12–14). This is largely due to its unique pharmacological profile, which supports its potential as an ideal target for migraine treatment, including lower risk for MOH and abuse liability, and absence of physical dependence (15). In addition, preclinical evidence demonstrates that DOR agonists are strikingly effective in acute and chronic models of migraine-associated pain (16, 17), post-traumatic headache (PTH) (18), and MOH (16). Moreover, DOR agonists have shown beneficial effects in modulating cortical spreading depression (CSD) (12, 19), an electrophysiological correlate of migraine with aura (MWA), that underlies symptoms including visual, speech and auditory disturbances, tingling and numbness (20, 21).

Significant advances have been made in identifying the molecular mechanisms underlying DOR activation–mediated analgesic and anti-migraine effects. DOR is a Gi/o protein-coupled receptor (GPCR) expressed at both presynaptic terminals and postsynaptic membranes (22–24), and activation of DOR can significantly reduce neuronal excitability (25), via inhibition of adenylyl cyclase activity, and subsequent protein kinase A (PKA) signaling (26, 27). In addition, DOR activation stimulates opening of GIRK potassium channels, leading to membrane hyperpolarization, and inhibits calcium channels, thereby reducing presynaptic calcium influx and neurotransmitter release (26). Together, these intracellular mechanisms can modulate the transmission and amplification of pain signals in the trigeminal system and ultimately contribute to the anti-migraine effects of DOR agonists.

Despite the anti-migraine effects, and the lower risk for adverse effects compared to other strategies like mu-opioid receptor (MOR) agonists, some DOR agonists have been associated with convulsions and the development of analgesic tolerance following repeated administration (28–30). These adverse effects have been suggested to result, from receptor internalization, β-arrestin–dependent signaling, and subsequent receptor desensitization, which together may limit sustained therapeutic efficacy (29). Critically however, these effects are not intrinsic consequences of DOR activation *per se* but instead reflect ligand-specific signaling profile properties (30, 31). This phenomenon, known as functional selectivity or biased agonism, refers to the ability of different agonists acting at same receptor to produce different active receptor states, which in turn trigger differential effects at the receptor or intracellular level (32). For instance, the high level of DOR internalization induced by the agonist SNC80 has been associated with convulsions (28, 30, 33), whereas the low-internalizing agonist ARM390 shows non-convulsant effects (29, 30, 34).

Therefore, drug development efforts have shifted towards developing signaling-biased ligands or partial agonists that selectively engage distinct subsets of receptor-mediated pathways, to avoid convulsions and analgesic tolerance. PN6047 (3-[[4-(dimethylcarbamoyl)phenyl]-[1-(thiazol-5-ylmethyl)-4-piperidylidene]methyl]benzamide), a novel and selective G protein-biased DOR agonist structurally derived from SNC80, has shown effectiveness in producing a robust analgesic effect in neuropathic pain (sciatic nerve ligation), osteoarthritic pain, and inflammatory pain (carrageenan) models (35). Strikingly, chronic PN6047 treatment produces neither analgesic tolerance nor convulsant effects (35). In addition, PN6047 has completed Phase I clinical trial and showed excellent safety and tolerability. Although PN6047 has shown safe effects in models of pain (35), its efficacy in preclinical models of headache remains to be evaluated. In the current study, we aimed to examine the effects of PN6047 in blocking cephalic allodynia in nitroglycerin-induced chronic migraine-associated pain, PTH, and MOH; and determine if PN6047 induces MOH and blocks CSD events, and CSD-induced allodynia.

## Materials and methods

### Animals

All animal experiments were performed according to the Association for Assessment and Accreditation of Laboratory Animal Care (AAALAC) guidelines administered by the IACUC committee at Washington University in St. Louis. Both male and female C57BL6/J mice (8-week age, Jackson Laboratoriy, Bar Harbor, ME. USA) were used in all experiments. For the PTH model, only male mice were used. For cortical spreading depression (CSD) in freely behaving mice, CSD was optogenetically triggered by activation of channelrhodopsin (ChR) in transgenic male and female mice that expressed the ChR2-eYFP fusion protein in layer 5 cortical neurons (8-week age, Strain 7612–B6.Cg-Tg, Thy1-COP4/EYFP,18Gfng/J; Thy1/ChR2; Jackson Laboratory, Bar Harbor, ME, USA). The animals were housed in groups of 3-5 mice, except Thy1-mice which were housed individually during recording. All mice were maintained in a sterilized solid bottom cage with contact bedding under controlled temperature on a 12:12h light/dark cycle, with *ad libitum* access to food and water. All behavioral experiments occurred in the light cycle between 8am – 5pm.

### NTG-induced hypersensitivity

Nitroglycerin (NTG, 10mg/kg, i.p., American Regent, Shirley, NY) or vehicle was injected every other day for 9 days (1, 3, 5, 7, and 9) after measuring basal mechanical cephalic threshold. On day 10, mechanical threshold was measured before (basal responses), and 45min after injecting vehicle or PN6047 (3mg/kg, i.p., post-treatment responses).

### Model of post-traumatic headache (PTH)

Mild traumatic brain injury (mTBI) was induced by using the closed head weight-drop method, as was previously described (18). Briefly, male mice were anesthetized with 2% isoflurane with an oxygen flow rate of 0.6-0.8 liters per minute. Mice were placed chest down on a foam sponge (dimensions: 7-1/2 in. × 5-1/2 in. × 1-7/8 in) to support the head and body, which allowed for anterior-posterior motion without any rotational movement during the impact. The mouse and sponge were placed directly underneath the weight-drop device which consisted of a hollow cylindrical tube (inner diameter 2.54 cm, 80 cm height) placed approximately 1cm vertically over the mouse’s head, in between the ear and eye. To induce mTBI, a 30g weight (13 mm diameter, 34 mm height) was dropped through the tube, striking the mouse and causing a closed head injury. Following mTBI, mice were returned to their home cages for recovery for 2 weeks. To model PTH, a low dose of NTG (0.1mg/kg, i.p) or vehicle were injected every other day for 9 days (1, 3, 5, 7, and 9). Basal mechanical cephalic threshold was measured on day 1, 5, and 9 before the injection. On day 10, basal mechanical threshold was measured before, and 45min after injecting vehicle or PN6047 (3mg/kg, i.p).

### Medication-overuse Headache (MOH)

Mice received daily injections of vehicle or sumatriptan (0.6mg/kg, i.p) for 11 days, and 15min before each injection mice were injected with PN6047 (3mg/kg, i.p) or vehicle. Basal mechanical cephalic threshold was measured on days 1, 5 and 8 before the injections. Follow-up measurements were conducted on day 12 (about 18h following the last injection).

### Sensory sensitivity testing

For cephalic allodynia, mice were tested in 4 oz paper cups included in the testing apparatus during habituation. The threshold for response to punctate mechanical stimuli was assessed using the up and down method. The plantar surface of the hind paw or the periorbital region caudal to the eyes near the midline was tested with a series of eight von Frey filaments (bending force ranging from 0.008 to 2 g). For hind paw testing, a response was characterized as lifting or shaking of the paw following stimulation. For cephalic testing, a response was defined as a shaking or ducking of the head, or vigorous grooming of the periorbital region following stimulation.

### Cortical spreading depression (CSD) model

Cortical spreading depression was performed as previously described (12, 36). Briefly, mice were anesthetized with isoflurane (induction 2–3 %; maintenance 0.75-1.25%; in 67% N2/ 33% O2) and placed in a stereotaxic frame on a homeothermic heating pad. Core temperature (37.0 ± 0.5°C), oxygen saturation (∼90%), heart rate, and respiratory rate (30–120 bpm) were continuously monitored during the experiment (PhysioSuite; Kent Scientific Instruments, Torrington, CT, USA). Mice were repeatedly tested with hind paw pinch to ensure that proper anesthetic levels were maintained. Following induction, the scalp was opened, a small area of the skull was thinned to make a cortical window (∼0.5 mm from sagittal, and ∼ 1.4 from coronal and lambdoid sutures), and mineral oil was applied to allow for greater visualization. For video recording a green LED (520 nm) light illuminated the skull throughout the experiment (1-UP; LED Supply, Randolph, VT, USA). Reflectance was collected with a lens (HR Plan Apo 0.5 × WD 136) through a 515LP emission filter on a Nikon SMZ 1500 stereomicroscope (Nikon Instruments, Melville, NY, USA). Images were acquired at 1.75 Hz using a high-sensitivity USB monochrome CCD (CCE-B013-U; Mightex, Pleasanton, CA, USA) with 4.65-micron square pixels and 1392 × 1040 pixel resolution. Two burr holes were drilled lateral to the thinned window around the midpoint of the rectangle. The burr holes were drilled deeper than the thinned skull portion such that the dura was exposed, but not so deep that the dura was broken. In the caudal burr hole, a pulled glass pipette filled with saline was inserted to record local field potentials (LFP). The electrode was connected to a ground wire placed underneath the skin, caudal to the skull, which was then connected to an amplifier. Following insertion of the electrode, an hour of background activity was recorded and monitored. This allowed stabilization if a CSD occurred during surgery. Baselines were recorded for an hour after which a glass pipette filled with 1M KCl was placed into the rostral burr hole. An initial drop of KCl was formed and then the flow of KCl was maintained. After 400s from the initial KCl drop, mice were injected with vehicle, SNC80 (5mg/kg, i.p) or PN6407 (3mg/kg, i.p), and recorded for 1h. Mice that showed at least 2 CSDs in the first 400s were included in the analysis. Four mice were excluded for not meeting this criteria. Video and subsequent LFP recordings were used to confirm CSD counts.

### Microchip system

#### Chip Implantation

For long term continuous recordings in freely behaving animals, microchips were permanently affixed to the skull in a short (<20 min) surgical procedure as described previously(37, 38). Briefly, mice were anesthetized in an induction chamber with isoflurane (3%). Once deeply anesthetized and not responding to toe pinch, they were placed in a stereotaxic frame where anesthesia was maintained using isoflurane (2%). Mice were kept on a heating pad during the procedure. Following disinfection of the scalp, the skull was surgically exposed, and a microchip was glued onto the skull with cyanoacrylate and permanently enclosed with black two-component resin (J-B Kwik, J-B Weld Company, TX, USA). Carprofen (2mg/kg) tablets were given for at least 2 days following surgery. No evidence of significant distress or pain was observed 1 week following surgery.

#### CSD triggering and recording

The chip system comprised an optical sensor to measure reflection, green LEDs for reflectance/optical intrinsic signal (OIS illumination), a blue LED for optical stimulation of ChR2, and a movement sensor(38). All chips were embedded on a small circuit board which also contained a waterproof connector so that the animals could be disconnected and reconnected if needed. The weight of the system, including the encapsulating resin, is less than 1 gram. OIS and movement data in these long-term experiments were continuously sampled with 128 Hz. CSD threshold was determined in two steps: First, general CSD was triggered optically by administering sequential pulses of light with the blue LED of increasing duration (first duration 1s) until CSD was observed based on OIS signal. After each light pulse, OIS was monitored for 5 min for the occurrence of CSD. If CSD occurred, no further pulses were delivered, whereas if no CSD occurred, then the duration was increased with 1s with the subsequent pulse. The general threshold represents an approximate range in seconds rather than an exact value in milliseconds. In the second step, a specific CSD threshold was determined by applying light pulses within a narrower range around the general threshold. For example, if the general threshold was 5.0s, the specific threshold test started at 4.0s and increased in 0.1s increments up to 5.0s, with a 5-minute interval between pulses. CSD occurrence was again verified based on the OIS signal. For each animal, CSD threshold was determined at the baseline, and approximately after 10 min of SNC80 (5 mg/kg, i.p), PN6047 (3 mg/kg, i.p) or vehicle injection, with ∼20hours between each session.

#### CSD triggering and sensory test

CSD was triggered after measuring basal mechanical threshold. After 1h of CSD, mice were injected with PN6047 (3mg/kg, i.p) or vehicle, and mechanical threshold was tested 1h post-drug treatment. The CSD was triggered optically by administering suprathreshold a single pulse of light for 10s. CSD was confirmed based on OIS signal. CSD occurred in all animals tested. Animals were included in the test only if CSD occurred following simulation. No mice were excluded.

### Data analysis

GraphPad Prism (version 10 and 11, GraphPad, San Diego, CA, USA) was used to perform the statistical analyses. For chronic administration experiments, two or three-way repeated measures ANOVA were used. Significant main effects and interactions were further pursued using Holm-Sidak’s post hoc by test only when a significant interaction was observed. All data are expressed as Mean ± Standard Error of the Mean (SEM). The accepted significance value for all tests was set at *P*<0.05. Detailed statistical analysis can be found in the Supplementary Statistics Table 1.

## Result

### Chronic NTG-induced cephalic allodynia is blocked by PN6047 treatment

To examine the anti-migraine effects of PN6047 in a migraine-associated pain model, mice were injected with vehicle or the human migraine trigger, NTG (10mg/kg, i.p), every other day for 9 days (Fig. 1A). Cephalic mechanical responses were measured on days 1, 5 and 9 before injections (Basal responses, Fig. 1A). Over time, a significant reduction in cephalic mechanical thresholds was observed in the NTG group compared to the vehicle group (Fig. 1B).

**Figure 1.**
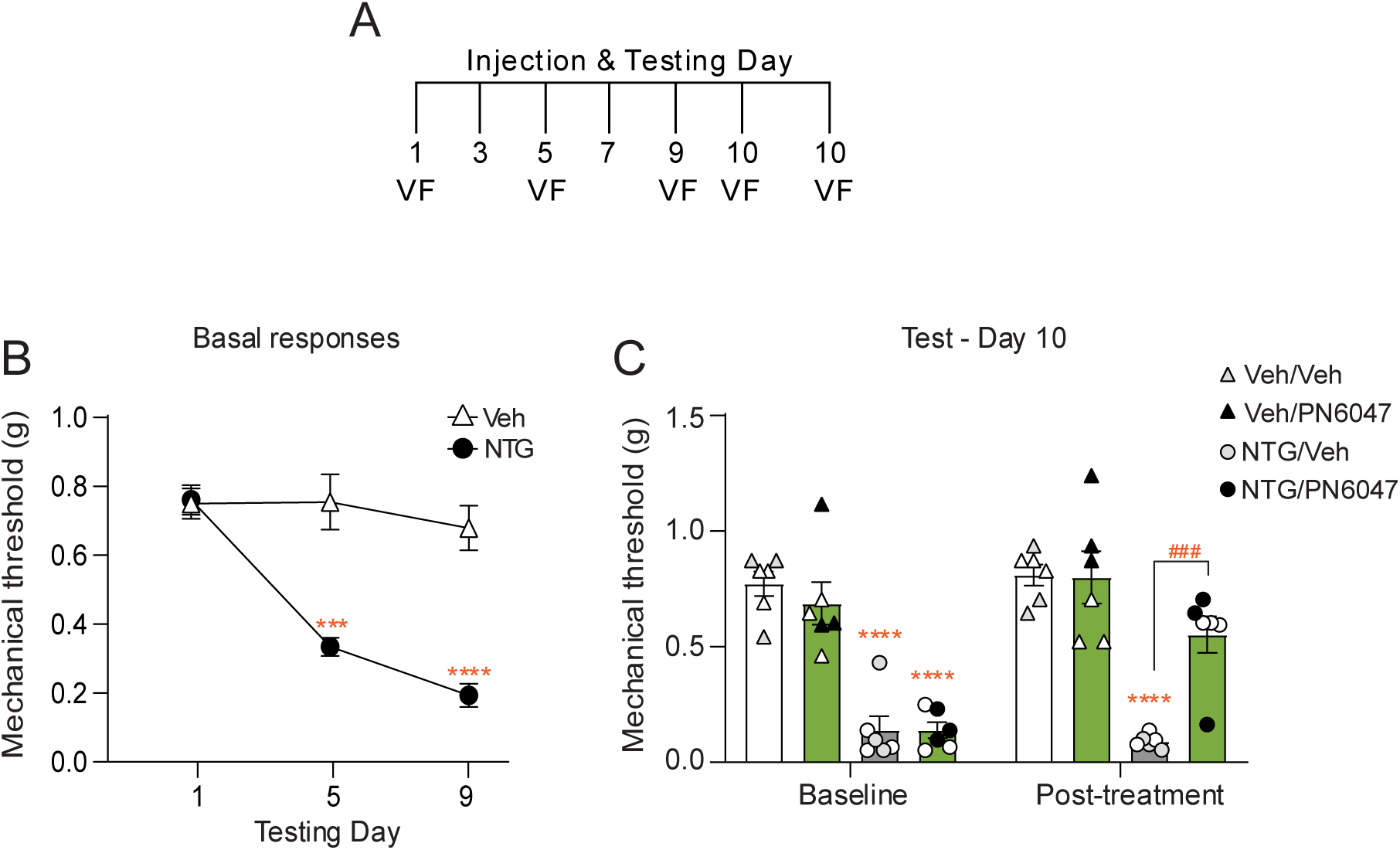
Cephalic allodynia induced by chronic intermittent NTG is blocked by PN6047 treatment. **(A)** The experimental timeline, mice were injected with vehicle or NTG (10 mg/kg, i.p) every other day for 9 days. Baseline of cephalic mechanical threshold was measured before Veh/NTG injection on day 1, 5, and 9. On day 10, baseline responses were determined, and mice were injected with vehicle or PN6047 (3 mg kg, i.p). Following 45min of the injection post-treatment responses were determined. **(B)** Baseline of cephalic mechanical threshold before Veh/NTG treatment on day 1, 5, and 9. Two-way RM ANOVA (*P* < 0.0001 time, dose, interaction), Holm–Sidak post hoc analysis, \*\*\**P* < 0.0001, \*\*\*\**P* < 0.0001, significantly different from vehicle group at the same time-point, n = 12 per group. **(C)** On day 10, basal cephalic threshold was determined, and mice were injected with vehicle or PN6047 (3 mg kg, i.p) and tested 45min later. PN6047 significantly reversed mechanical allodynia induced by chronic NTG treatment. Three-way repeated measures ANOVA (*P* < 0.05 time–treatment–drug interaction), Holm–Sidak post hoc analysis, \*\*\*\**P* < 0.0001, compared to vehicle/vehicle group at the same time point, ^###^*P* < 0.001, significantly different from NTG/vehicle group at the same time point. n = 6 per group, results from female animals are indicated with white symbols. Means ± SEM.

We next determined if chronic allodynia induced by NTG was blocked by PN6047, on day 10 (∼20h after last Veh/NTG injection). Baseline mechanical responses were determined, and mice treated with NTG continued to show significant allodynia (Fig. 1C, baseline). Mice were injected with vehicle or PN6047 (3mg/kg, i.p), and tested 45min later (post-treatment responses, Fig. 1A). PN6047 treatment significantly blocked NTG-induced cephalic allodynia (Fig. 1C, post-treatment) and did not affect general nociception in mice chronically treated with vehicle (Fig. 1C, Veh-PN6047). These results indicate that established cephalic allodynia induced by chronic treatment with NTG can be blocked by acute PN6047 treatment.

### Acute treatment with PN6047 blocks post-traumatic headache

We have previously developed a mouse model of post-traumatic headache (PTH) where mice receive a mTBI and show hyposensitivity to low dose NTG (18). Here, mice underwent a closed head injury followed by a 2-week recovery period (Fig. 2A). At 2 weeks post-injury, low dose NTG (0.1 mg/kg, i.p) or vehicle was administered every other day over 9 days (5 days total, Fig. 2A). Basal cephalic mechanical responses were measured before mTBI, and on days 1, 5 and 9 before Veh/NTG treatment (Fig. 2B).

**Figure 2.**
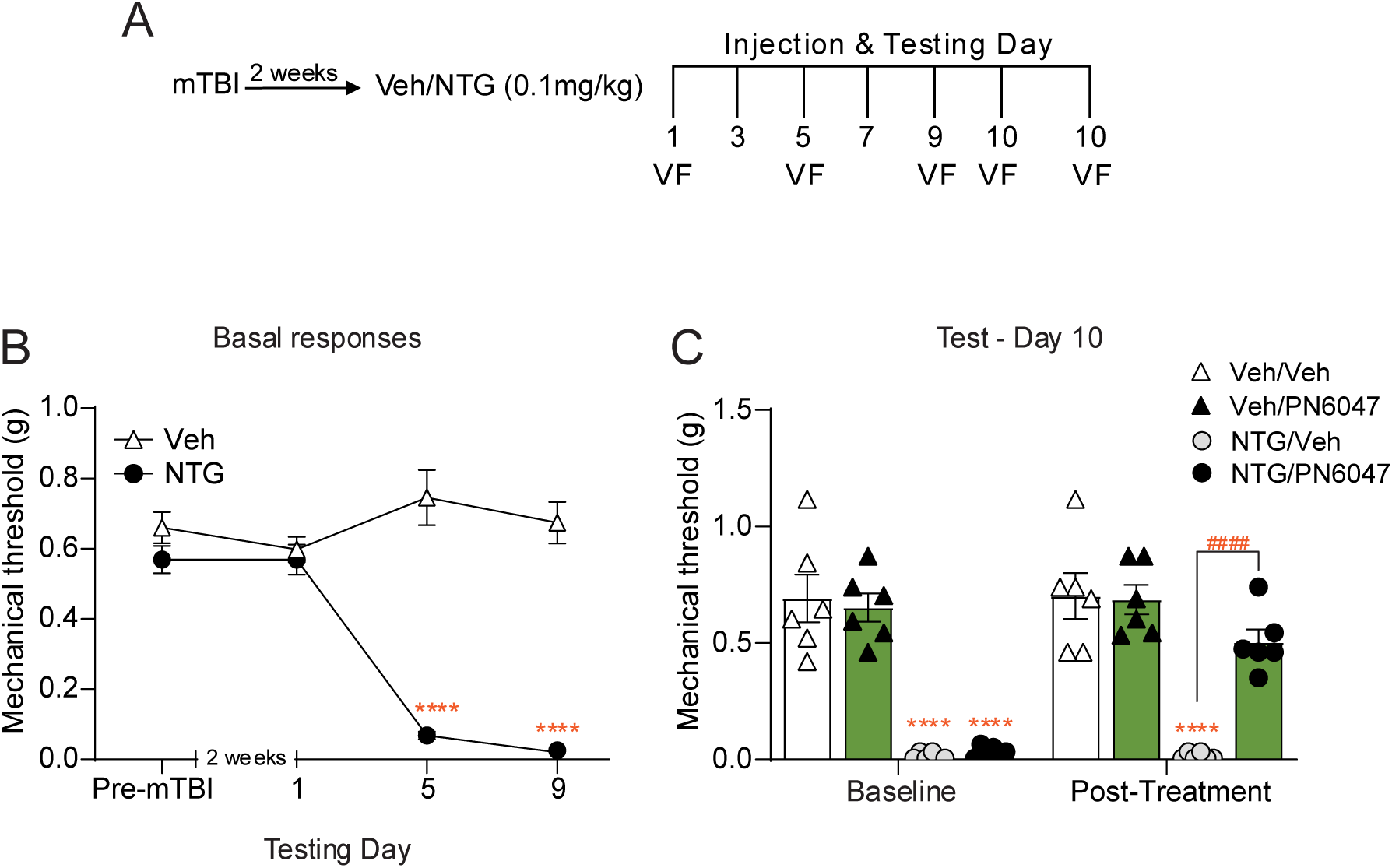
PN6047 treatment blocks cephalic allodynia in a model of PTH. **(A)** Experimental timeline, cephalic mechanical threshold was measured before mTBI. All mice underwent a closed head weight drop (mTBI), and 2 weeks later were treated with vehicle or low dose NTG (0.1 mg/kg, i.p) every other day for 9 days. Baseline cephalic mechanical responses were measured before Veh/NTG injection on day 1, 5, and 9. On day 10, baseline responses were determined, and mice were injected with vehicle or PN6047 (3 mg/kg, i.p). Following 45min of the injection post-treatment responses were determined. **(B)** Baseline cephalic mechanical threshold before mTBI, and Veh/NTG treatment. No difference was observed in cephalic mechanical threshold before mTBI, and 2 weeks post-mTBI on Veh/NTG treatment day 1. Low dose of NTG (0.1 mg/kg, i.p) produced significant allodynia following mTBI. Two-way repeated measures ANOVA (*P* < 0.0001 time, dose, interaction), Holm–Sidak post hoc analysis, \*\*\*\**P* < 0.0001, significantly different from vehicle at the same time point, n = 12 per group. **(C)** On day 10, baseline cephalic threshold was determined, and mice were injected with vehicle or PN6047 (3 mg/kg, i.p) and tested 45min later. PN6047 significantly blocked established PTH-associated allodynia. Three-way repeated measures ANOVA (*P* < 0.05 time–treatment–drug interaction), Holm–Sidak post hoc analysis. \*\*\*\**P* < 0.0001, compared to Veh/Veh group at the same time point, ^####^*P* < 0.001, significantly different from NTG/Veh group at the same time point. n = 6 per group. Means ± SEM. Only males were used in this experiment due to weight restrictions for mTBI induction.

No difference was observed in cephalic mechanical threshold before mTBI (Fig. 2B, Pre-mTBI), and 2 weeks post-injury (Fig. 2B, day 1). We observed a significant reduction in cephalic mechanical thresholds in the mTBI/NTG treatment group compared to the mTBI/vehicle group on days 5, and 9 (Fig. 2B), this cephalic allodynia was still present ∼18h following the final NTG injection (Day 9, Fig. 2B). On day 10, baseline responses were determined, and mice were injected with PN6047 (3mg/kg, i.p) or vehicle, and cephalic responses were determined 45min later (Fig. 2C). PN6047 treatment significantly blocked cephalic allodynia induced by mTBI+chronic low dose of NTG (Fig. 2C). Notably, PN6047 treatment showed no effect on the vehicle treatment group (Fig. 2C). This data demonstrates that DOR activation by acute treatment of PN6047 can alleviate established PTH-associated pain.

### PN6047 treatment inhibits sumatriptan-induced medication overuse headache

Chronic use of acute migraine medications such as triptans are associated with medication overuse headache (MOH) and considered as a risk factor for migraine chronification (8, 9). Here, we tested whether chronic PN6047 could inhibit sumatriptan (SUMA)-induced MOH, and if PN6047 itself causes MOH. To model MOH, SUMA (0.6mg/kg, i.p) or vehicle, in combination with vehicle or PN6047 (3mg/kg, i.p) were injected once daily for 11 days (Fig. 3A). Basal cephalic mechanical thresholds were measured before each injection on days 1, 5, and 8 (Fig. 3A). Mice chronically treated with SUMA/Veh showed a significant reduction in cephalic mechanical threshold (Fig. 3B), and this allodynia was still present ∼18h following the final SUMA injection (Day 12, Fig. 3B).

**Figure 3.**
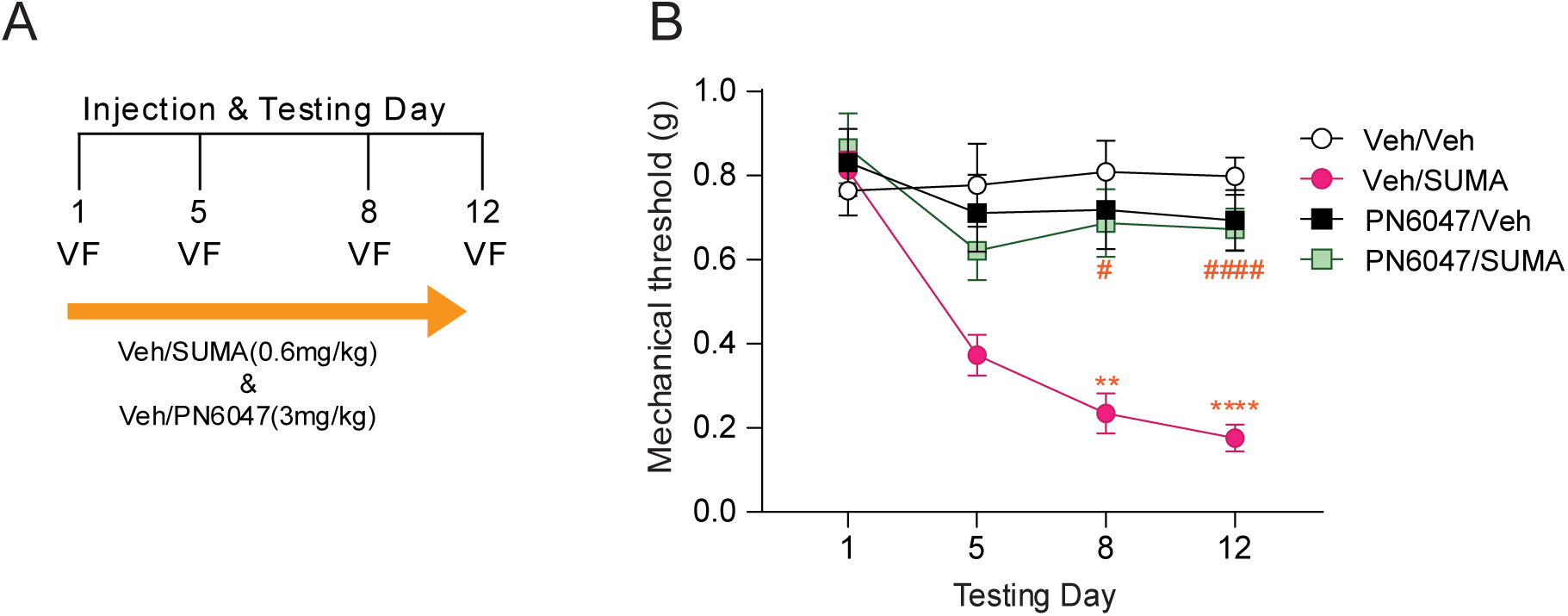
Chronic PN6047 prevents development of MOH and does not cause MOH. **(A)** Experimental timeline, mice were co-administered vehicle or PN6047 (3mg/kg, i.p), with vehicle or sumatriptan (SUMA, 0.6mg/kg, i.p) every day for 11 days. Baseline cephalic mechanical responses were measured before injection on day 1, 5, 8, and 12 (∼20h after the final injection). **(B)** Chronic PN6047 treatment significantly prevented sumatriptan-induced chronic allodynia. Three-way repeated measures ANOVA (*P* < 0.05 time–treatment–drug interaction), Holm–Sidak post hoc analysis, \*\**P* < 0.01, \*\*\*\**P* < 0.0001 between vehicle/SUMA group and vehicle/vehicle group at the same time point, ^#^*P* < 0.05, ^####^*P* < 0.0001 between vehicle/SUMA group and PN6047/SUMA group at the same time point, n = 12 per group. Means ± SEM.

Strikingly, chronic treatment of PN6047 significantly prevented the development of cephalic allodynia induced by chronic treatment with SUMA (Fig. 3B). More importantly, chronic treatment of PN6047 showed no significant effect on cephalic mechanical threshold (Fig. 3B, PN6047-Veh). The current data demonstrates that chronic PN6047 does not cause MOH and can prevent the development of MOH caused by sumatriptan overuse, without tolerance.

### PN6047 reduces and modulates behaviors associated with cortical spreading depression (CSD)

The ability of drugs to reduce the number of CSD events has been successfully used as a method for predicting the effects of migraine preventive medication (39, 40). Previously we demonstrated that DOR agonists including SNC80 and KNT127 reduced the number of CSD events in anesthetized mice (19, 41). Here, we examined whether PN6047 could reduce CSD events similar to the well-established DOR agonist SNC80 (19, 41). CSD events were triggered in anesthetized mice by applying a constant amount of KCl onto the dura while optical intrinsic signals (OIS) and local field potentials (LFP) were recorded. SNC80 (5mg/kg, i.p), PN6047 (3mg/kg, i.p) or vehicle were injected 400s after initial KCl application and CSD events were recorded for 1h following administration. The results demonstrate that both SNC80 and PN6047 treatments significantly decreased the number of CSD events compared to vehicle treatment (Fig. 4A, B). Notably, no significant difference was observed between SNC80 and PN6047 treatment in reducing the number of CSD events (Fig. 4A, B). These data indicate that PN6047, similar to other DOR agonists, has a beneficial effect in modulating CSD, an electrophysiological correlate of migraine with aura (MWA).

**Figure 4.**
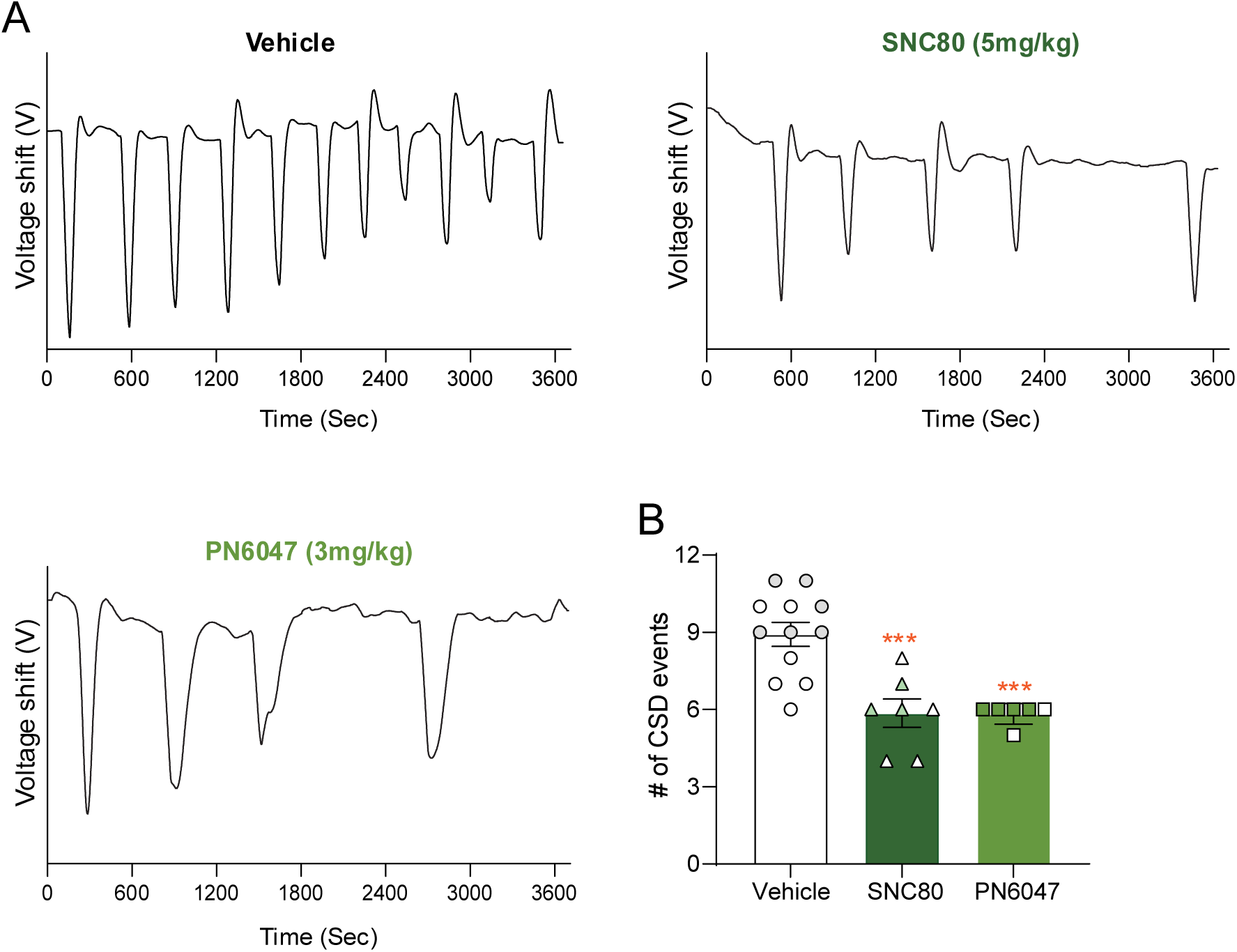
PN6047 exhibits comparable efficacy to SNC80 in reducing the number of CSD events in anesthetized mice. **(A)** Representative line tracing of CSD events over 1h period following injection of vehicle, SNC80 (5 mg/kg, i.p), or PN6047 (3 mg/kg, i.p). **(B)** Mice treated with the DOR agonists, SNC80 or PN6047, had a significant reduction in the average number of CSD events recorded at 1h compared to vehicle group. One-way ANOVA (*P* < 0.001 treatment). Holm–Sidak post hoc analysis. \*\**P* < 0.01, \*\*\**P* < 0.0001, compared to vehicle group. n = 6/12 per group, results from female animals are indicated with white symbols. Means ± SEM.

### DOR agonists increase the threshold to trigger CSD in a freely behaving mice along with subsequent allodynia

We wanted to confirm the beneficial effects of PN6047 in reducing CSD events in freely behaving mice using a long-term continuous recording chip (42). In this case, CSD was induced using optogenetic stimulation of the cortex in mice expressing channel rhodopsin (Thy-Chr2). First, individual CSD threshold was determined by applying blue light pulses of increasing duration (100ms) at 5-minute intervals between pulses. CSD occurrence was verified based on the OIS signal. For each animal, CSD threshold was determined at the baseline, and after approximately 10 min of SNC80 (5mg/kg, i.p), PN6047 (3mg/kg, i.p) or vehicle injection, with ∼20 hours between each session. Treatment with SNC80 or PN6047 significantly increased CSD threshold by increasing the duration of the light activation needed to induce CSD (Fig. 5A), with thresholds increasing by 40% and 31%, respectively, whereas vehicle treatment did not alter the threshold relative to baseline (Supplementary Fig. 1A).

**Figure 5.**
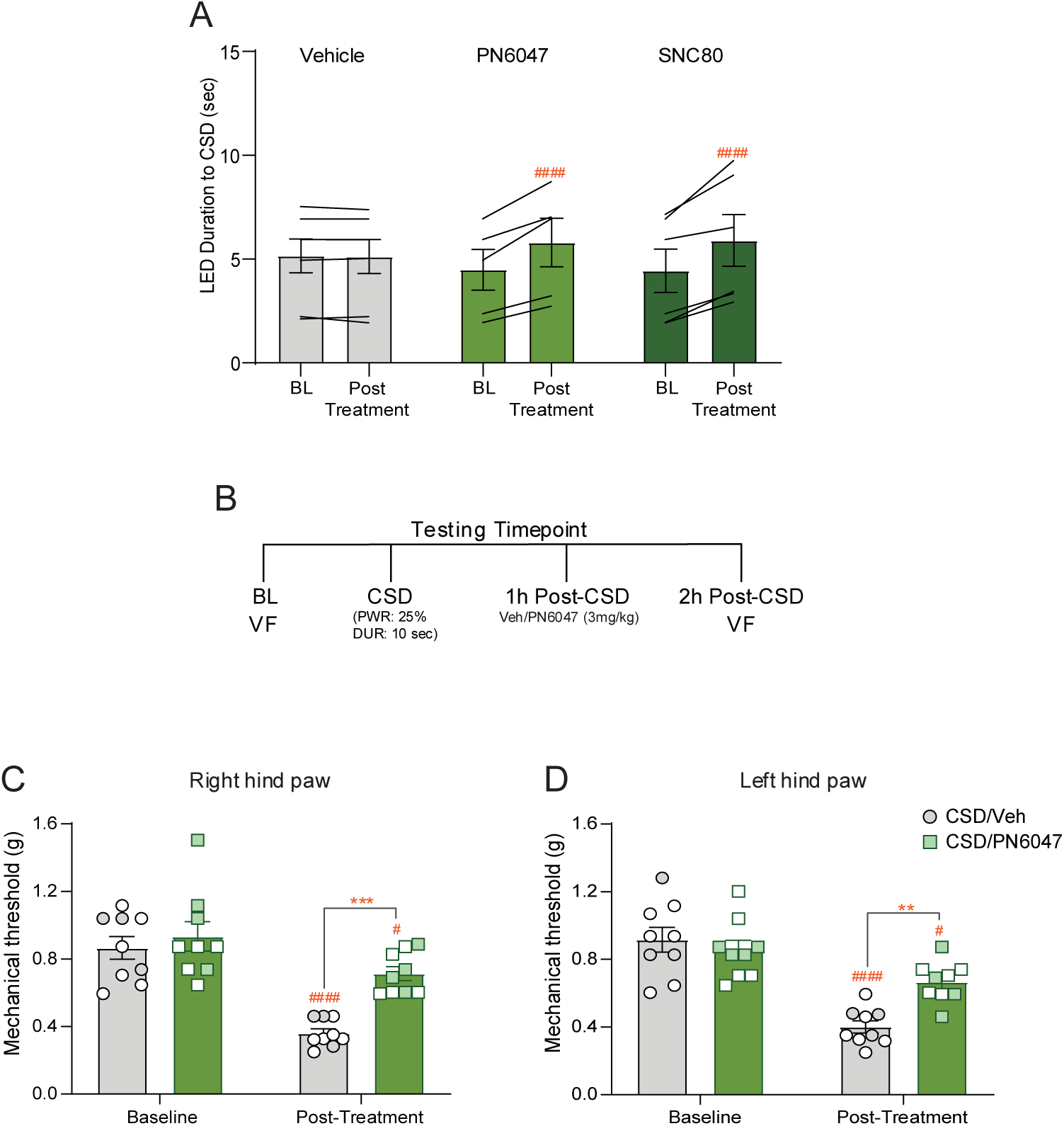
DOR agonists, SNC80, and PN6047 delay triggered CSD in a freely behaving mouse along with CSD-evoked allodynia. **(A)** CSD was optically triggered by blue light stimulation in a freely behaving Thy1Chr mouse implanted with a microchip on the skull, at baseline (BL) and after injection of vehicle or DOR agonist. Treatment with the DOR agonist increased the LED activation threshold (longer activation duration), indicating a stronger stimulation is needed to trigger CSD, i.e. a higher threshold. Two-way repeated measures ANOVA (*P* < 0.05 time–treatment–interaction). Holm–Sidak post hoc analysis. ###*P* < 0.0001, compared to the baseline of the same group, n = 5-7 per group. **(B)** Experimental timeline, peripheral (right/left hind paw) mechanical responses were measured before CSD. The CSD was triggered optically by administering suprathreshold single pulse of light for 10s. CSD was confirmed based on OIS signal. After 1h of CSD, mice were injected with vehicle or PN6047 (3 mg kg, i.p), and post-treatment responses were determined 1h later. **(C)** Baseline of right and **(D)** left hind paw before and 2h after CSD. PN6047 significantly inhibited CSD-induced peripheral allodynia compared to vehicle treatment. Two-way repeated measures ANOVA (*P* < 0.05 time–treatment–interaction). Holm–Sidak post hoc analysis. #*P* < 0.05, ####*P* < 0.0001, compared to the baseline of the same group, \*\**P* < 0.01, \*\*\**P* < 0.001, significant difference between CSD/Veh vs. CSD/PN6047 group. n = 9 per group, results from female animals are indicated with white symbols. Means ± SEM.

CSD is a well-established model of MWA (20), and previous evidence indicates that CSD can lead to cephalic and peripheral allodynia development in mice (43–45). Here, we examined if PN6047 treatment could block peripheral allodynia-induced by CSD. Peripheral mechanical baselines were measured prior to triggering CSD (Basal response, Fig. 5B), and 1h after injection of PN6047 (3 mg/kg, i.p) or vehicle (2h post-CSD response, Fig. 5B). CSD was triggered and confirmed using OIS signal. We found that a single triggered CSD event in an awake freely-behaving mouse produced significant mechanical allodynia in the right and left hind paws at 2h post-CSD which resolved by 24h post-CSD (Supplementary Fig. 1B). PN6047 treatment significantly blocked this mechanical allodynia, compared to vehicle treatment (Fig.5C, B). These data demonstrate that DOR activation by acute treatment with PN6047 can alleviate peripheral mechanical hypersensitivity induced by CSD.

## Discussion

In the current study, we examined the efficacy of PN6047, a novel G protein-biased DOR agonist, in blocking headache-associated pain in models of– NTG, CSD, PTH and MOH. Here, we found that PN6047 blocks chronic cephalic allodynia established by chronic intermittent NTG, and PTH. More importantly, chronic PN6047 treatment does not produce MOH and was effective in preventing the development of SUMA-induced MOH. In addition, PN6047 showed beneficial effects in reducing CSD events along with subsequent allodynia. Together, the current data demonstrate that PN6047 has potent analgesic and anti-migraine efficacy in preclinical models of migraine-associated pain, does not induce MOH or analgesic tolerance, and exhibits a favorable safety and tolerability profile.

Despite substantial advances in migraine treatment, a significant proportion of migraine patients remain dissatisfied with their medication due to limited efficacy, and adverse effects (10, 11). Importantly, patients suffering from PTH face similar limitations and the lack of specific pharmacological treatments for this disorder further contributes to the complexity of clinical management (46, 47). We have previously shown that experimental DOR agonists (SNC80 and KNT127) can block the development of chronic migraine (48) and PTH (18), as well as block established chronic allodynia associated with these disorders (18, 19). In this study we confirmed that PN6047, a DOR agonist that has undergone human phase I clinical trial, produces similar effects. Our data showed that PN6047 significantly blocks cephalic allodynia in chronic NTG-induce migraine-associated pain, and PTH. The specific molecular mechanisms underlying the anti-migraine effects of DOR activation in NTG- and PTH models may involve suppression of the endogenous migraine trigger calcitonin gene–related peptide (CGRP) signaling in the trigeminovascular system (22). The current findings further support that DOR is an effective and mechanistically distinct therapeutic target that could expand the treatment options for migraine and headache disorders.

Overuse of medication is considered a risk factor for MOH (8, 9), contributing to the cycle of headache chronicity. The primary treatment for MOH is cessation of the overused medication; nevertheless, clinical data indicate a 20-50% relapse rate within the first year of withdrawal, with most patients relapsing within the first 6 months of withdrawal (49–53). We had previously tested SNC80 in a model of SUMA-induced MOH and found that acute treatment relieved chronic allodynia established in this model. In the present study, we took advantage of the finding that, unlike other DOR agonists, PN6047 does not produce tolerance after chronic daily treatment (35), and we determined that concurrent treatment with SUMA could prevent the development of MOH. In the same experiment we also found that chronic treatment with PN6047 alone did not itself produce MOH, which is in contrast to chronic treatment with SNC80, which was reported to produce limited MOH in a similar paradigm (16). This difference may reflect ligand-specific signaling properties of PN6047 at DOR unlike those of SNC80 which promotes receptor internalization and subsequent desensitization, thereby limiting its therapeutic efficacy with repeated treatment (29), PN6047 has not been shown to induce these desensitizing effects (35). This property may contribute to its sustained therapeutic efficacy without developing analgesic tolerance or promoting MOH.

Interestingly, the protective effects of PN6047 in inhibiting MOH development were not due to an acute analgesic effect, as cephalic allodynia was measured prior to drug treatment. Instead, this effect may be related to the regulation of central sensitization induced by sumatriptan. Previous studies have shown that prolonged sumatriptan treatment increases markers of neuroinflammation and long-term synaptic plasticity including cFOS, and CGRP in the trigeminovascular system (54, 55). While there is currently no direct evidence demonstrating that DOR regulates these changes in MOH models, prior work indicates that DOR agonists can modulate neuroinflammation in preclinical models of Alzheimer’s disease (56), and downregulate CGRP levels in model of chronic migraine–associated pain (22). Together, these findings demonstrate that PN6047 prevents the development of MOH without itself inducing it, highlighting its favorable safety profile and therapeutic potential in migraine and headache disorders. Importantly, this protective effect is maintained during prolonged treatment, with no evidence of analgesic tolerance, suggesting that DOR activation by PN6047 acts as a homeostatic regulator that prevents the transition to a sensitized MOH state.

The high comorbidity between headache disorders and psychiatric conditions including emotional distress, anxiety, and depression probably contributes to headache chronicity and disability (57), and may also play a role in relapse (58). DOR agonists exert dual effects, as DOR activation can prevent hyperalgesia and positively regulate emotional tone. Accordingly, SNC80 and PN6047 have been reported to produce anxiolytic-like effects and anti-depressant-like behaviors in animals (35, 59). This further supports the potential clinical value of PN6047, as patients with MOH require support to manage hyperalgesia during overuse of medication, withdrawal, emotional stress, and the prolonged sensitivity that can persist for months (57, 58, 60, 61).

Approximately 30% of migraine patients experience aura as part of their migraine attacks (62). CSD is a slowly propagating wave of depolarization that is followed by inhibition of brain activity (20) and it is thought to be the electrophysiological phenomenon that underlies migraine aura (21). Our previous studies show that DOR agonists can reduce CSD events (19, 41), and confer an advantage over other acute migraine treatments such as triptans or opioids which have been shown to enhance CSD with repeated use (63, 64). Here, we demonstrated that the clinical compound, PN6047, significantly inhibited CSD events similar to the experimental DOR agonists SNC80 and KNT127. Furthermore, we also tested PN6047 and the comparator, SNC80, in a CSD model in freely behaving mice (42). We further demonstrate that PN6047 robustly delays CSD events in this novel optogenetic model, consistent with findings from the KCl-induced CSD model under anesthesia. The inhibitory effects of DOR agonists on CSD are potentially mediated by reducing neuronal excitability, which stabilizes network activity and disrupts the synaptic processes required for wave initiation and propagation. Future studies will explore these mechanisms further.

Clinical evidence indicates that headaches typically emerge approximately 1h after the onset of aura (65). Consistent with this, preclinical studies have shown that CSD induced by KCl application or optogenetic stimulation produces cephalic and peripheral mechanical allodynia (44, 66, 67). However, these approaches generally rely on invasive procedures, such as viral injection, cannula implantation, or are conducted under anesthesia. In contrast, a major advantage of the microchip approach is the ability to reliably evoke CSD in awake freely behaving animals using a minimally invasive, transcranial optogenetic stimulation paradigm (38). Our results demonstrate that a single CSD event induces peripheral mechanical allodynia within 2h, which was resolved 24h post-induction (Supplementary Fig. 1B). Notably, despite unilateral CSD induction, no differences were observed between ipsilateral and contralateral mechanical thresholds. Interestingly, even after CSD induction we found that PN6047 treatment significantly blocked peripheral allodynia induced by CSD, indicating that DOR agonists are not only effective at preventing CSD events, but can also be beneficial once CSD has been evoked. These results suggest that CSD induced central sensitization which contributes to peripheral allodynia, could be effectively blocked by DOR agonist treatment. The neuronal mechanisms underlying CSD-induced headache and peripheral allodynia may be related to evidence that CSD produces marked alterations in the cerebrospinal fluid (CSF) proteome, including a robust elevation in CGRP levels, a key mediator of trigeminal activation (68). Importantly, following CSD, the CSF was shown to flow from the subarachnoid space into the TG, providing a direct route by which centrally released proteins can access primary afferent neurons (68). This CNS–PNS communication pathway may facilitate the activation and sensitization of trigeminal nociceptors, thereby linking cortical events to peripheral sensitization and ultimately contributing to headache generation following CSD. DOR expression is observed all along this pathway, including in the dura (69), cortex, trigeminal ganglia, and trigeminal nucleus caudalis (22, 70), thus allowing for modulation by DOR agonists.

Our findings further support that DOR-based treatments are effective in models of migraine and headache disorders. Here, we demonstrated that PN6047, a novel G protein-biased DOR agonist that has recently completed Phase I clinical trial, with no indications of safety or tolerability concerns, blocks migraine-associated pain in chronic NTG-induced allodynia, and PTH models. Moreover, PN6047 shows high efficacy in reducing CSD events along with subsequent allodynia. Importantly, we found that chronic PN6047 treatment prevents the development of cephalic allodynia in a sumatriptan-induced MOH model, without inducing tolerance and MOH itself. Together, these data demonstrate that PN6047 has potent analgesic efficacy in preclinical models of headache-associated pain, does not induce MOH, and exhibits a favorable safety and tolerability profile. Overall, the data presented here supports the future clinical testing of PN6047 in headache disorders.

## Declarations

### Ethics approval and consent to participate

All animal experiments were performed according to the Association for Assessment and Accreditation of Laboratory Animal Care (AAALAC) guidelines administered by the IACUC committee at Washington University in St. Louis.

### Consent for publication

Not applicable.

### Availability of data and materials

The datasets generated during and/or analyzed during the current study are available from the corresponding author on reasonable request.

### Author contribution

Y.A.I. contributed to study design, data collection, data analysis, data interpretation, and drafting initial manuscript. Y.Z., contributed to data collection. S.M.B., M.A., G.C.F. and A.C. contributed to data interpretation, and technical support. E.J., J.T., D.K., and B.V.M., contributed to study design and manuscript editing. A.A.P. contributed to study design, data analysis, data interpretation, project administration, funding acquisition, and editing final manuscript version.

### Clinical trial number

not applicable.

## Acknowledgements

The authors would like to acknowledge the sudden loss of Dr. Emily Jutkiewicz during the course of this study. Dr. Jutkiewicz was a valued colleague and leader in the field of opioid receptor pharmacology and her loss is deeply felt.

