## Supplemental Figure 1 for "PN6047 Demonstrates Broad Preclinical Efficacy in Headache Models as a Novel Delta-Opioid Receptor Agonist"

**Supplementary Figure**

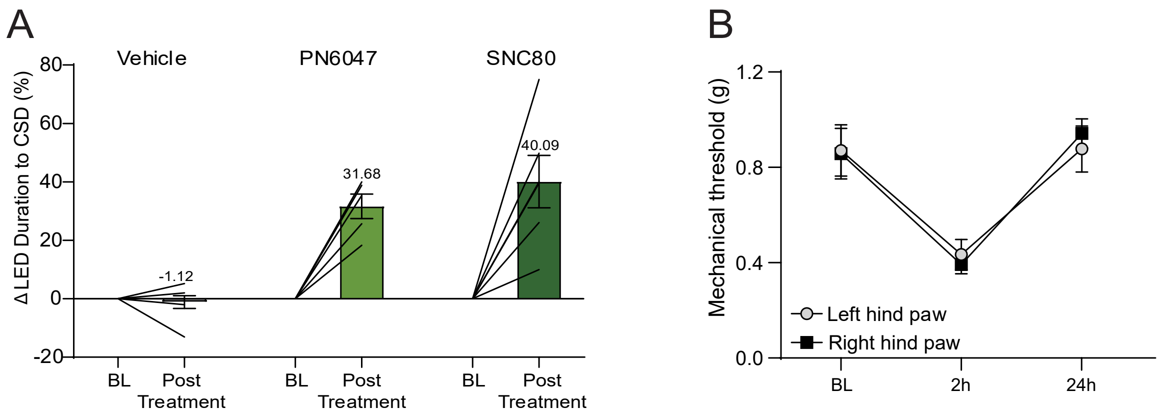


**Supplementary Figure 1: DOR agonists, SNC80, and PN6047 delay triggered CSD in a freely behaving mouse, and triggered CSD evokes short-term allodynia. (A)** CSD was optically triggered by blue light stimulation in a freely behaving Thy1Chr mouse implanted with a microchip on the skull, at baseline (BL) and after injection of vehicle or DOR agonist. Treatment with SNC80 or PN6047 increased CSD threshold by increasing the duration of the light activation needed to induce CSD, with thresholds increasing by 40% and 31%, respectively, whereas vehicle treatment did not alter the threshold relative to baseline. n = 5-7 per group. **(B)** Peripheral mechanical responses were measured before CSD (BL). The CSD was triggered optically by administering suprathreshold single pulse of light for 10s. CSD was confirmed based on OIS signal. Post-treatment responses were determined 2h and 24 later. A single CSD event induces peripheral mechanical allodynia observed at 2h and resolved by 24h post-induction. 2-way RM ANOVA $p<0.001$ time. n = 4 per group. Means ± SEM.
